# MODERATE ALCOHOL CONSUMPTION INDUCES LASTING IMPACTS ON PREFRONTAL CORTICAL SIGNALING IN MICE

**DOI:** 10.1101/2024.04.03.587955

**Authors:** Grace C. Smith, Keith R. Griffith, Avery R. Sicher, Dakota F. Brockway, Elizabeth A. Proctor, Nicole A. Crowley

## Abstract

Both alcohol use disorder (AUD) and Alzheimer’s Disease and Related Dementias (ADRD) appear to include disruption in the balance of excitation and inhibition in the cortex, but their potential interactions are unclear. We examined the effect of moderate voluntary binge alcohol consumption on the aged, pre-disease neuronal environment by measuring intrinsic excitability and spontaneous neurotransmission on prefrontal cortical pyramidal (excitatory, glutamatergic) and non-pyramidal (inhibitory, GABAergic) neurons following a prolonged period of abstinence from alcohol in mice. Results highlight that binge alcohol consumption has lasting impacts on the electrophysiological properties of prefrontal cortical neurons. A profound increase in excitatory events onto layer 2/3 non-pyramidal neurons following alcohol consumption was seen, along with altered intrinsic excitability of pyramidal neurons, which could have a range of effects on Alzheimer’s Disease progression in humans. These results indicate that moderate voluntary alcohol influences the pre-disease environment in aging and highlight the need for further mechanistic investigation into this risk factor.

**HIGHLIGHTS:**

- Moderate voluntary alcohol consumption leads to disruption of prefrontal cortical signaling at protracted ages in both male and female mice
- Excitatory glutamatergic events onto prelimbic cortex GABAergic neurons, but not pyramidal neurons, are increased 6 months following alcohol exposure
- Both pyramidal neurons and GABAergic neurons show altered intrinsic excitability 6 months following alcohol exposure

## 1. INTRODUCTION

Healthy neuronal functioning and cell signaling in the brain requires a delicate balance of excitatory and inhibitory activity across neural circuits. Disruption to this balance occurs independently in both Alzheimer’s Disease (AD) and with excessive alcohol use, but the impact of alcohol use on the progression of excitation and inhibition in the aging brain is unknown. The impact of alcohol on AD is heavily debated, where some studies find alcohol use worsens cognitive decline and others find mild alcohol consumption may improve cognition (Piazza-Gardner et al., 2013; Peng et al., 2020; Seemiller et al., 2024). Due in part to the conflicting results on the potentially protective effect of alcohol against cognitive decline, the World Health Organization recommends against the consumption of alcohol (WHO, 2019).

Recently, investigations into the underlying neuronal activity in patients diagnosed with AD have shown network abnormalities and interneuron dysfunction in cortical-hippocampal circuits early in disease progression (Palop & Mucke, 2016; Ramírez-Toraño et al., 2021). Maintaining balance between excitatory and inhibitory (E/I) inputs is a multidimensional process that includes neuronal homeostasis and changes in gene and/or protein expression (Sohal & Rubenstein, 2019). Imbalance in E/I has been hypothesized to play a role in AD, potentially due to interneuron dysfunction (i.e., dysregulation in inhibition), failure of homeostatic regulation of firing, and hyperactivity in cortico-hippocampal circuits (Styr & Slutsky, 2018). Innervation of the cortex by the hippocampus is critical to the functioning of the medial prefrontal cortex (O’Donnell & Grace, 1995). The cortico-hippocampal circuit is formed by ipsilateral, unidirectional, glutamatergic, monosynaptic projections from the CA1 and subicular ventral hippocampus onto pyramidal and non-pyramidal neurons in layer 2/3 of the medial prefrontal cortex (Jay et al., 1992; Parent et al., 2010). Lesion of the medial prefrontal cortex results in cognitive deficits that resemble AD symptoms (Kolb, 1991). Additionally, layer 2/3 of the cortex has been implicated in maintaining optimal cortical E/I balance (Shew et al., 2011), and the prefrontal cortex is implicated in a plethora of brain disorders (Brockway & Crowley, 2020) - overall indicating the importance of cortico-hippocampal circuits in AD-related cognitive deficits.

Alcohol consumption is likely one environmental mechanism for chronic E/I imbalance; however, the effects of alcohol on AD are inconsistent - some studies report increased risk of AD with alcohol use, others report no effect, and some even find a protective effect of low dose alcohol (Piazza-Gardner et al., 2013). These conflicting results highlight the need for mechanistic and causative investigations into the relationship between alcohol and AD. Acute alcohol consumption is known to alter the normal E/I balance through region and cell-type specific changes in excitation and inhibition in the cortex. In the mouse prelimbic (PL) cortex, exposure to voluntary alcohol decreases the excitability of both somatostatin- and vasoactive intestinal peptide-expressing interneurons, resulting in disinhibition of pyramidal neurons and an increase in pyramidal intrinsic excitability (Dao et al., 2021; Thompson et al., 2023), while exposure to chronic ethanol alters the action potential threshold in pyramidal neurons (Pleil et al., 2015). However, despite the plethora of evidence that disruptions in E/I balance occur immediately following alcohol consumption, it remains unknown how long these changes in excitability persist following acute alcohol consumption. Here, we sought to understand the effect of voluntary alcohol consumption via the Drinking-in-the-Dark (DID) binge like model of ethanol on measurements of intrinsic excitability and synaptic transmission in the prefrontal cortex approximately six months after the cessation of alcohol in wildtype mice.

## 2. MATERIALS AND METHODS

### 2.1 Mice

Experiments detailed in this paper used 50 mice; all experiments were approved by the Pennsylvania State University Institutional Animal Care and Use Committee. Male and female (25 M, 25 F) C57BL/6J (stock #000664, The Jackson Laboratory) mice were bred at Jackson Laboratory and delivered at 7 weeks of age. At 9 weeks of age, mice were placed into single housing on a 12 h reversed light cycle (lights off at 07:00 am). At 8 weeks of age, the drinking-in-the-dark (DID) paradigm of voluntary alcohol consumption was conducted as previously published (Rhodes et al., 2005; Thiele & Navarro, 2014; Dao et al., 2021; Crowley et al., 2019).

### 2.2 Drinking-In-The-Dark

Mice received 20% (v/v) ethanol (EtOH, Koptec, Decon Labs, King of Prussia, PA) diluted in tap water for 2 h beginning 3 h into the dark cycle (i.e., 10 am – 12 pm) for three consecutive days. On the fourth day (binge day) mice received EtOH for 4 h (i.e., 10 am – 2 pm), followed by three days of water only. This was repeated for a total of four weeks. Mice had *ad libitum* access to food and water aside from the 2 or 4 h period the water bottle was replaced with an EtOH sipper tube. Following the four weeks of DID, mice remained single housed and undisturbed until 9-12 months of age, with *ad libitum* access to food and water.

### 2.3 Electrophysiology

Electrophysiology was conducted between 9-12 months of age (+/- 2 weeks; corresponding to about 6 months following the cessation of alcohol consumption) similar to our previously published work (Brockway et al., 2023; Sicher et al., 2023). Mice were exposed to an isoflurane chamber for deep anesthetization, the brain was quickly extracted and processed according to the N-methyl-D-glucamine (NMDG) protective recovery method (Ting et al., 2014). Immediately following extraction, brains were placed into ice-cold oxygenated NMDG-HEPES artificial cerebrospinal fluid (aCSF) for one minute (in mM: 92 NMDG, 2.5 KCl, 1.25 NaH_2_PO_4_, 30 NaHCO_3_, 20 HEPES, 25 glucose, 2 thiourea, 5 Na-ascorbate, 3 Na-pyruvate, 0.5 CaCl_2_•2H_2_O, 10 MgSO_4_•7H_2_O; pH 7.3-7.4). The brain was then blocked for sectioning of the PL cortex. Using a Compresstome vibrating microtome VF-300-0Z (Precessionary Instruments, Greenville, NC), the PL cortex (as identified by the Allen Mouse Brain Atlas) was sectioned into 300 µM coronal slices. Slices were then transferred to heated (31 °C) oxygenated (95% O_2_, 5% CO_2_) NMDG-HEPES aCSF for 8 min, then to heated oxygenated normal aCSF (containing the following, in mM: 124 NaCl, 4.4 KCl, 2 CaCl_2_, 1.2 MgSO_4_, 1 NaH_2_PO_4_, 10.0 glucose, 26.0 NaHCO_3_; pH 7.4, mOsm 305-310), where they were allowed to recover for at least 1 h before electrophysiological experiments. Glass recording electrodes were pulled from thin-walled borosilicate glass capillary tubes to 3-6 MΩ with a Narshige P-100 Puller (Narshige International USA, Amityville, NY).

Whole-cell voltage clamp and current clamp recordings were performed in pyramidal and non-pyramidal (putative GABAergic) neurons in layer 2/3 of the PL cortex. Neurons were identified by previously published categorizations of morphology (triangular soma and apical dendrites for pyramidal, smaller round soma for non-pyramidal), and membrane characteristics (pyramidal: capacitance > 75 pF, or membrane resistance < 100 MΩ) (Dao et al., 2021). Neurons recorded were determined to be healthy for further experimental investigations based on the following criteria: access resistance < 40 MΩ, resting membrane potential lower than –40 mV, higher than –90 mV.

Intrinsic excitability experiments were recorded with a potassium gluconate intracellular recording solution (K-Gluc, in mM: 135 K-Gluc, 5 NaCl, 2 MgCl_2_, 10 HEPES, 0.6 EGTA, 4 Na_2_ATP, 0.4 Na_2_GTP; pH 7.35, mOsm 287-290). The resting membrane potential (RMP), rheobase (minimum current required to elicit an action potential), action potential (AP) threshold (the membrane potential when the first action potential fired), fast afterhyperpolarization amplitude (fAHP, the difference between the AP threshold and the lowest membrane potential immediately following the AP), AP half-width (the time in ms at the midpoint between the AP threshold and AP peak), and current injection (VI) induced firing were assessed. Rheobase was measured using an increasing ramp of current injection until the first AP. VI-induced firing was assessed using 10 pA stepwise increase in applied current from 0-200 pA. Experiments were conducted at both the resting membrane potential, then at the common holding potential of –70 mV.

Spontaneous excitatory and inhibitory postsynaptic currents (sEPSC and sIPSC, respectively) were recorded with a cesium-methanesulfonate intracellular recording solution (Cs-Meth, in mM: 135 Cs-methanesulfonate, 10 KCl, 10 HEPES, 1 MgCl_2_, 0.2 EGTA, 4 MG-ATP, 0.3 GTP, 20 phosphocreatine; pH 7.33, mOsm 287-290) with 1 mg Lidocaine. Spontaneous events were recorded in gapfree, at a holding potential of – 55 mV and +10 mV for sEPSCs and sIPSCs, respectively, for 10 min. Only the last two minutes were analyzed.

### 2.4 Analysis

Recorded signals were digitized at 10 kHz and filtered at 3 kHz using a Multiclamp 700B amplifier and analyzed using Clampfit 11.2.1 software (Molecular Devices, Sunnyvale, CA) and custom Matlab code (Smith et al., 2024). Custom Matlab code was written to analyze the current injection induced firing, acquiring the number of action potentials at each step of current injection. Spontaneous postsynaptic currents were analyzed using Clampfit event detection, where a template was made for each recording and the 2 min file was analyzed matching events to the template; from this analysis measures of frequency (Hz) and average amplitude (pA) were recorded. For each measure, a maximum of three neurons per subtype, per mouse were recorded.

Whenever possible, experimenter was blinded to cell-type, alcohol condition, and sex. Initial statistical analysis was performed using GraphPad Prism 10.1.2, and outliers were identified and excluded using ROUT tests (Q = 1%). Data were analyzed by two-way analysis of variance (ANOVA) and post-hoc testing where indicated, with a significance threshold of α = 0.05. Q-Q plots were made to ensure normality of data for ANOVA. Data presented below indicates means and standard error of the mean (SEM). Additionally, correlations between each measure and the animal’s total EtOH consumption were performed.

### 2.5 Partial Least Squares Multivariate Modeling

We used partial least squares analysis (Geladi & Kowalski, 1986), a supervised statistical modeling tool to account for co-variation in related electrophysiological excitability parameters. Briefly, partial least squares analysis identifies an optimal set of eigenvectors to relate the matrix of observed parameters (here, electrical properties) to the matrix of phenotypic observations with minimum residual error. Models were constructed using PLS_Toolbox (Eigenvector Research) in MATLAB. Data was normalized along each parameter by Z-score. Cross-validation was performed with one-third of the dataset. The number of latent variables was chosen to minimize cumulative error over all predictions. In cases where two phenotypic groups are being discriminated or one continuous variable is being regressed against, we have orthogonalized our model to maximize the discrimination or regression to the first latent variable. All reported accuracies and root-mean-square-error are from cross-validation unless otherwise noted. We calculated model confidence by randomly permuting the phenotype (outcome) matrix, while leaving the matrix of observed parameters intact to preserve the topography of the parameter space. The cross-validated accuracies of these 100 random models form a distribution to which we compared the accuracy of our model using a Mann-Whitney U test. Importance of each parameter to the model predictive accuracy was quantified by VIP score, an average of the weights of each parameter over all latent variables normalized by latent variable percent of variance explained. VIP score greater than 1 denote an above-average contribution and was considered important for model performance and prediction.

## 3 RESULTS

### 3.1 Basal electrophysiological properties in PFC neurons do not differ across sex or alcohol exposure

Male and female mice that underwent DID consumed EtOH to binge levels during the four-hour exposure day; consistent with previously published work on sex differences in alcohol consumption, female mice consume more overall EtOH over the full four week exposure (two-tailed t-test, t(30) = 2.281, p = 0.0298; overall model in Fig. **1A**, EtOH consumption in Fig. **1B**). Basal membrane characteristics for each cell type (pyramidal and non-pyramidal, GABAergic) were compared, and were largely not significantly different based on DID condition or sex. Pyramidal membrane capacitance (pF) was not significantly different across sex or drinking condition (F_sex_(1, 58) = 0.1661, p = 0.6851; F_DID_(1, 58) = 1.688, p = 0.1990; F_sex x DID_ (1, 58) = 3.592, p = 0.0631; Fig. **1C**). An interaction effect of sex and DID was present in pyramidal membrane resistance (MΩ), with post-hoc testing revealing a significantly lower resistance in female control mice compared to male control mice (F_sex_(1, 58) = 3.256, p = 0.0764; F_DID_(1, 58) = 0.9779, p = 0.3268; F_sex x DID_ (1, 58) = 7.100, p = 0.0100; Fig. **1D**), but no alterations from DID. There was no significant difference in non-pyramidal membrane capacitance or resistance across sex or DID (capacitance: F_sex_(1,48) = 0.5619, p = 0.4571; F_DID_(1, 48) = 1.749, p = 0.1922; F_sex x DID_(1, 48) = 0.6716, p = 0.6719; Fig. **1E**; resistance: F_sex_(1,47) = 1.102, p = 0.2991; F_DID_(1, 47) = 2.155, p = 0.1487; F_sex x DID_(1, 47) = 1.130, p = 0.2932; Fig. **1F**).

**Figure 1.**
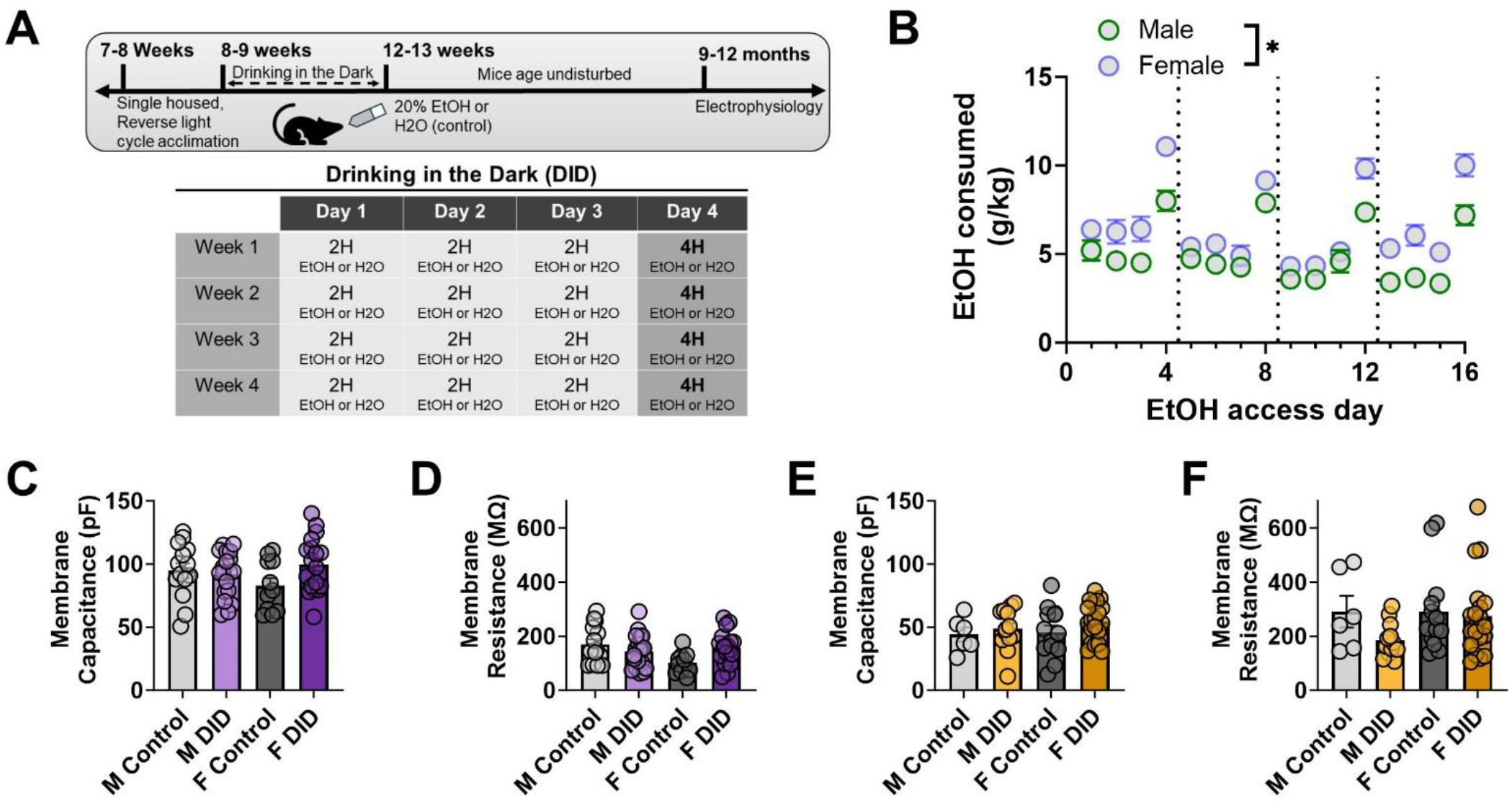
Experimental timeline and comparison of PL cortical neuron basal electrophysiological characteristics. **A)** Experimental timeline. Mice received 4 weeks of exposure to the voluntary binge-alcohol consumption model ‘drinking in the dark’ (DID) during early adulthood. They then remained undisturbed for approximately six months until further experiments. **B)** Average EtOH consumption (g/kg) for each day of DID across males and females. **C)** Pyramidal membrane capacitance (pF) was not significantly different across groups. **D)** Pyramidal membrane resistance (MΩ) significantly differs in a sex by drinking manner, where male control neurons had higher membrane resistance than female control neurons. **E)** Non-pyramidal membrane capacitance (pA) was not significantly different. **F)** Non-pyramidal membrane resistance (MΩ) was not significantly different. Error bars indicate SEM.

### 3.2 Modest alcohol exposure induces long-lasting adaptations in cortical pyramidal neuron intrinsic excitability in both male and female mice

Assessment of DID-induced changes in intrinsic excitability of pyramidal neurons following a prolonged period of abstinence from alcohol was conducted to determine if DID has lasting effects on PL cortical signaling. Intrinsic excitability measurements were conducted on a total of 62 pyramidal neurons from the following groups: female control: n = 11 cells/6 mice, female DID: n = 18 cells/9 mice, male control: n = 14 cells/5 mice, male DID: n = 19 cells/8 mice.

Pyramidal neurons in mice that were exposed to the DID paradigm of both sexes had hyperpolarized RMP compared to mice that were not exposed to DID (F_sex_(1, 58) = 3.252, p = 0.0765; F_DID_(1, 58) = 4.198, p = 0.0450; F_sex x DID_ (1, 58) = 0.4371, p = 0.5112; Fig. **2B**). There was a significant decrease amongst the DID condition in AP threshold in pyramidal neurons at resting membrane potential (F_sex_(1, 58) = 0.02049, p = 0.8867; F_DID_(1, 58) = 4.431, p = 0.0396; F_sex x DID_ (1, 58) = 0.08178, p = 0.7759; Fig. **2C**). The rheobase was not significantly different across drinking condition at resting membrane potential but was significantly different across sex, with males showing an average 114.8 pA rheobase (+/- 13.42 SEM), and females showing an average 146.0 pA rheobase (+/- 16.98 SEM) (F_sex_(1, 54) = 4.100, p = 0.0478; F_DID_(1, 54) = 0.8581, p = 0.3584; F_sex x DID_ (1, 54) = 2.761, p = 0.1024; Fig. **2D**). No significant difference in fAHP was observed across sex or DID (F_sex_(1, 56) = 0.2124, p = 0.6467; F_DID_(1, 56) = 0.007532, p = 0.9311; F_sex x DID_ (1, 56) = 3.289, p = 0.0751; Fig. **2E**). The AP half-width of pyramidal neurons was not significantly different across groups (F_sex_(1, 58) = 0.7545, p = 0.3886; F_DID_(1, 58) = 0.8725, p = 0.3541; F_sex x DID_ (1, 58) = 3.808, p = 0.0558; Fig. **2F**). There was no difference in AP threshold while holding the cell at –70 mV (F_sex_(1, 55) = 0.01702, p = 0.8967; F_DID_(1, 55) = 1.629, p = 0.2073; F_sex x DID_ (1, 55) = 0.2641, p = 0.6094; Fig. **2G**). The rheobase was not significantly different across drinking condition and sex at the common holding potential of –70 mV (F_sex_(1, 58) = 3.630, p = 0.0617; F_DID_(1, 58) = 0.2477 p = 0.6206; F_sex x DID_ (1, 58) = 1.136, p = 0.2910; Fig. **2H**).

**Figure 2.**
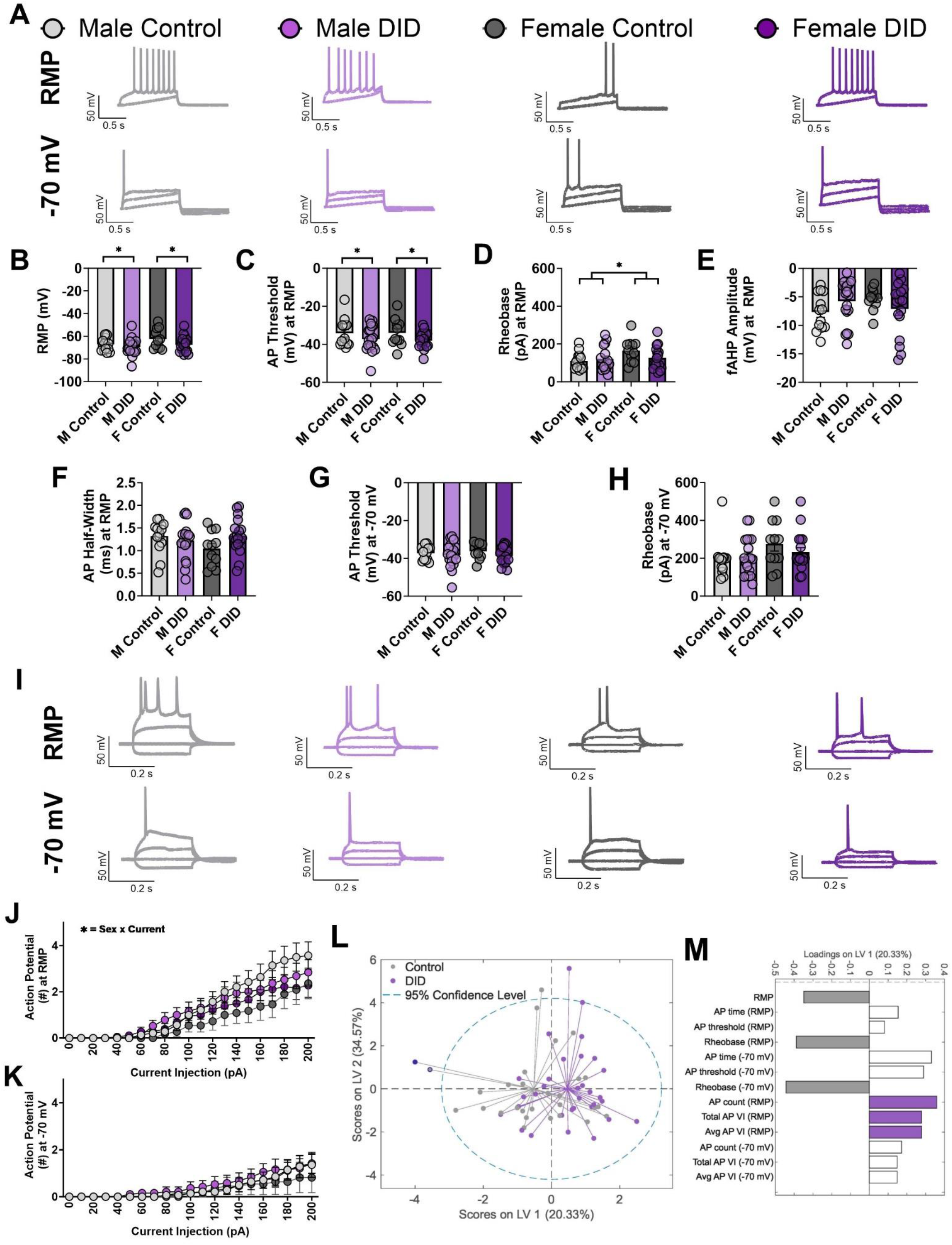
PL cortex pyramidal neuron intrinsic excitability is altered following voluntary alcohol consumption and extended abstinence across both sexes. **A)** Representative traces of rheobase at resting membrane potential (top row) and –70 mV (bottom row). **B)** RMP (mV) is significantly hyperpolarized in DID mice compared to control mice. **C)** AP threshold (mV) at RMP is significantly more negative in DID mice compared to control mice. **D)** Rheobase (pA) is significantly higher female mice compared to male mice. **E)** Ampitude of fAHP is not significantly different across groups. **F)** No significant differences in AP half-width (ms) at RMP. **G)** AP threshold while holding the cell at –70 mV is not significantly different across groups. **H)** Rheobase at –70 mV is not significantly different. **I)** Representative traces of VI, showing current injection of –100, 0, 100, and 200 pA at RMP (top row) and –70 mV (bottom row). **J)** The number of action potentials fired in VI at RMP is significantly different in a sex by current dependent manner. **K)** The number of action potentials fired in VI at –70 mV is not significantly different across groups. **L)** Partial least squares scores plot on the first two latent variables demonstrates separation in overall excitability of pyramidal neurons in control vs. DID mice. Model is orthogonalized to maximize the projection of separation onto the first latent variable. **M)** Loadings on the first latent variable, which contains the majority of differentiation between control vs. DID pyramidal neurons. Positive loadings indicate increases in DID pyramidal neurons, negative loadings indicate decreases in DID pyramidal neurons. Colored bars represent parameters with VIP score > 1, indicating greater than average contribution to predictive accuracy.

The number of action potentials fired in response to the VI protocol was analyzed using a three-way ANOVA across sex, DID, and current injection step with a significant interaction between current and sex at RMP (F_current_(1.427, 82.75) = 60.77, p <0.0001; F_sex_(1, 58) = 2.007, p = 0.1619; F_DID_(1, 58) = 0.03275, p = 0.8570; F_current x sex_(20, 1160) = 2.189, p = 0.0019; F_current x DID_(20, 1160) = 0.6999, p = 0.8294; F_sex x DID_(1, 58) = 0.5243, p = 0.4719; F_current x sex x DID_(20, 1160) = 0.9031, p = 0.5834; Fig. **2J**). It is expected that current is a significant driver of AP number, and that was the only significant effect at –70 mV (F_current_ 1.397, 81.03) = 18.55, p < 0.0001; F_sex_ (1, 58) = 0.2963, p = 0.5883; F_DID_ (1, 58) = 0.3422, p = 0.5608; F_current x sex_ (20, 1160) = 0.2625, p = 0.9996; F_current x DID_ (20, 1160) = 0.2323, p = 0.9998; F_sex x DID_ (1, 58) = 0.007999, p = 0.9290; F_current x sex x DID_ (20, 1160) = 0.4499, p = 0.9825; Fig. **2K**). In summary, significant long-term adaptations in pyramidal intrinsic excitability following DID in middle aged mice included hyperpolarized resting membrane potential, and decreased AP threshold at RMP.

We recognize that these excitability parameters are related and partially co-dependent, and that univariate analysis may therefore not account for the co-variance between them. To verify that we are not losing any biologically important relationships in the patterns of the co-variation among these individual parameters, we constructed supervised multivariate statistical models that use excitability parameters to blindly predict animal phenotype. Using partial least squares analysis, we identified that pyramidal neurons in the DID group (regardless of quantity of alcohol consumption) displayed reduced resting membrane potential and action potential threshold, while increasing action potential time (Fig. **2L, M**: 6 latent variables, 63% accuracy, 73% confidence). The difference in action potential threshold was strongest at resting membrane potential, while the difference in action potential time was strongest at –70 mV.

### 3.3 Non-Pyramidal (putatively GABAergic) intrinsic excitability was not altered following alcohol consumption

The same experiments outlined above were also conducted on non-pyramidal cells in layer 2/3 of the PL cortex. Experiments for non-pyramidal neurons outlined below had the following cell counts-female control: n = 12 cells/6 mice, female DID: n = 20 cells/9 mice, male control: n = 8 cells/4 mice, male DID: n = 13 cells/7 mice. There were no significant differences in the resting membrane potential of non-pyramidal neurons (F_sex_(1,48) = 0.01860, p = 0.8921; F_DID_(1, 48) = 0.1253, p = 0.7249; F_sex x DID_(1, 48) = 0.08625, p = 0.7703; Fig. **3B**). No significant differences in AP threshold were observed at RMP (F_sex_(1,49) = 0.2394, p = 0.6268; F_DID_(1, 49) = 0.9695 p = 0.3296; F_sex x DID_(1, 49) = 0.04104, p = 0.8403; Fig. **3C**). No significant differences in the rheobase at RMP were observed (F_sex_(1, 46) = 0.7470, p = 0.3919; F_DID_(1, 46) = 0.3482, p = 0.5580; F_sex x DID_(1, 46) = 0.1146, p = 0.7365; Fig. **3D**). The amplitude of fAHP was not significantly different in terms of sex or DID (F_sex_(1, 45) = 1.178, p = 0.2836; F_DID_(1, 45) = 1.855, p = 0.1800; F_sex x DID_(1, 45) = 0.26877, p = 0.6068; Fig. **3E**). No significant differences were observed in AP half-width in non-pyramidal neurons across sex or DID condition (F_sex_(1,48) = 0.1374, p = 0.7126; F_DID_(1, 48) = 1.671, p = 0.2023; F_sex x DID_(1, 48) = 0.1891, p = 0.6656; Fig. **3G**). In both sexes, DID groups had a significant increase in AP threshold at –70 mV (F_sex_(1,47) = 0.8990, p = 0.3479; F_DID_(1, 47) = 12.82, p = 0.0008; F_sex x DID_(1, 47) = 1.925, p = 0.1718; Fig. **3G**). There were no significant differences in rheobase at –70 mV holding potential (F_sex_(1,49) = 0.1908, p = 0.6642; F_DID_(1, 49) = 1.321, p = 0.2560; F_sex x DID_(1, 49) = 0.1404, p = 0.7095; Fig. **3H**).

**Figure 3.**
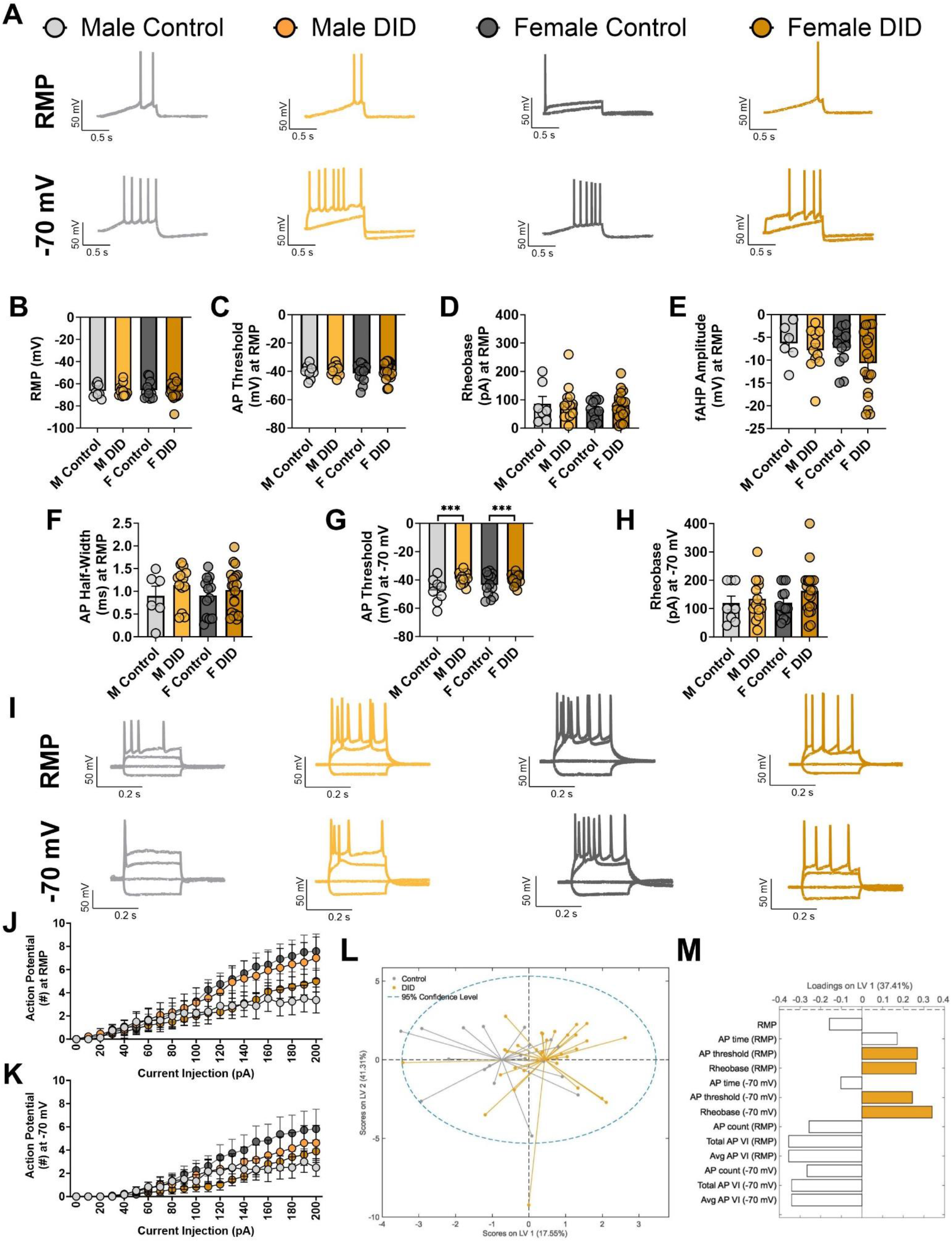
PL cortex non-pyramidal intrinsic excitability is largely unchanged following voluntary binge drinking and extended abstinence in both sexes. **A)** Representative traces of rheobase at resting membrane potential (top row) and –70 mV (bottom row). **B)** No significant difference in resting membrane potential across groups. **C)** No significant difference in AP threshold at RMP. **D)** No significant difference in rheobase at RMP. **E)** No significant difference in fAHP amplitude. **F)** AP half-width at RMP is not significantly different. **G)** A significant depolarization in AP threshold in DID mice compared to control mice. **H)** No change in rheobase at –70 mV. **I)** Representative traces of VI, showing current injection of –100, 0, 100, and 200 pA at resting membrane potential (top row) and –70 mV (bottom row). **J)** A current by sex by DID effect is present in the number of APs fired during VI at RMP. **K)** A current by sex by DID effect is present in the number of APs fired during VI at -70 mV. **L)** Partial least squares scores plot demonstrates separation in overall excitability of pyramidal neurons in control vs. DID mice. Second latent variable is shown for visualization purposes only and was not included in the final model for prediction or allocation of loadings. **M)** Loadings on the first latent variable. Positive loadings indicate increases in DID pyramidal neurons, negative loadings indicate decreases in DID pyramidal neurons. Colored bars represent parameters with VIP score > 1, indicating greater than average contribution to predictive accuracy.

A current by sex by DID effect was observed in the number of AP fired during VI at RMP (F_current_(1.821, 89.23) = 44.67, p <0.0001; F_sex_ (1, 49) = 0.06455, p =0.8005; F_DID_(1, 49) = 0.007573, p = 0.9310; F_current x sex_(20, 980) = 0.7163, p = 0.8121; F_current x DID_(20, 980) = 0.1903, p > 0.9999; F_sex x DID_(1, 49) = 3.013, p = 0.0889; F_current x sex x DID_(20, 980) = 4.234, p < 0.0001; Fig. **3J**) and at –70 mV (F_current_(1.583, 72.82) = 27.06, p <0.0001; F_sex_ (1, 46) = 0.09851, p =0.7550; F_DID_(1, 46) = 0.3068, p = 0.5823; F_current x sex_(20, 920) = 0.8158, p = 0.6959; F_current x DID_(20, 920) = 0.4410, p = 0.9844; F_sex x DID_(1, 46) = 1.559, p = 0.2182; F_current x sex x DID_(20, 920) = 1.724, p = 0.0250; Fig. **3K**). Post hoc Tukey’s test indicated that at 200 pA, female control neurons fire the most APs (mean = 7.583), male control neurons fire the least (mean = 3.375), with DID neurons in between (female mean = 5.000, male mean = 7.000).

We again performed multivariate analysis using partial least squares to account for any co-variance in electrophysiological parameters that may obscure underlying relationships or interfere with interpretation. In agreement with the univariate analysis, we could not significantly differentiate non-pyramidal excitability between DID and non-drinking mice (Fig. **3L, M**: 1 latent variable, 52% accuracy, 65% confidence).

### 3.4 Moderate alcohol consumption does not alter spontaneous post-synaptic currents onto pyramidal neurons

Spontaneous excitatory and inhibitory postsynaptic currents (sESPCs and sIPSCs, respectively) were recorded from pyramidal neurons by voltage-clamping the cells to –55 mV (sEPSC) and +10 mV (sIPSC) (example traces in Fig. **4A****, 4D** respectively). A total of 68 pyramidal neurons in layer 2/3 were recorded in the following groups: female control: n = 15/6 mice, female DID: n = 25/10 mice, male control: 9 cells/4 mice, male DID: 19 cells/9 mice. No significant differences in sEPSC (-55 mV) frequency across DID condition or sex were observed (F_sex_(1, 59) = 0.5617, p = 0.4566; F_DID_ (1, 59) = 0.7659, p =0 .3850; F_sex X DID_ (1, 59) = 0.4617, p = 0.4995; Fig. **4B**). No significant differences in amplitude were seen (F_sex_(1, 63) = 0.3205, p = 0.5733; F_DID_(1, 63) = 0.9437, p = 0.3350; F_sex x DID_(1, 63) = 0.7718, p = 0.3830; Fig. **4C**). Similarly, no changes across DID condition or sex were present in sIPSC (+10 mV) frequency (F_sex_(1, 64) = 0.1343, p = 0.7152; F_DID_(1, 64) = 1.538, p = 0.2194; F_sex x DID_(1, 64) = 1.884, p = 0.1747; Fig. **4E**) or sIPSC amplitude (F_sex_(1, 63) = 0.2426, p = 0.6241; F_DID_(1, 63) = 0.6818, p = 0.412; F_sex x DID_(1, 63) = 2.974, p = 0.0895; Fig. **4F**). No significant differences in synaptic drive were observed (F_sex_(1, 62) = 1.728, p = 0.1935; F_DID_(1, 62) = 0.2988, p = 0.5866; F_sex x DID_(1, 62) = 1.100, p = 0.2983; Fig. **4G**).

**Figure 4.**
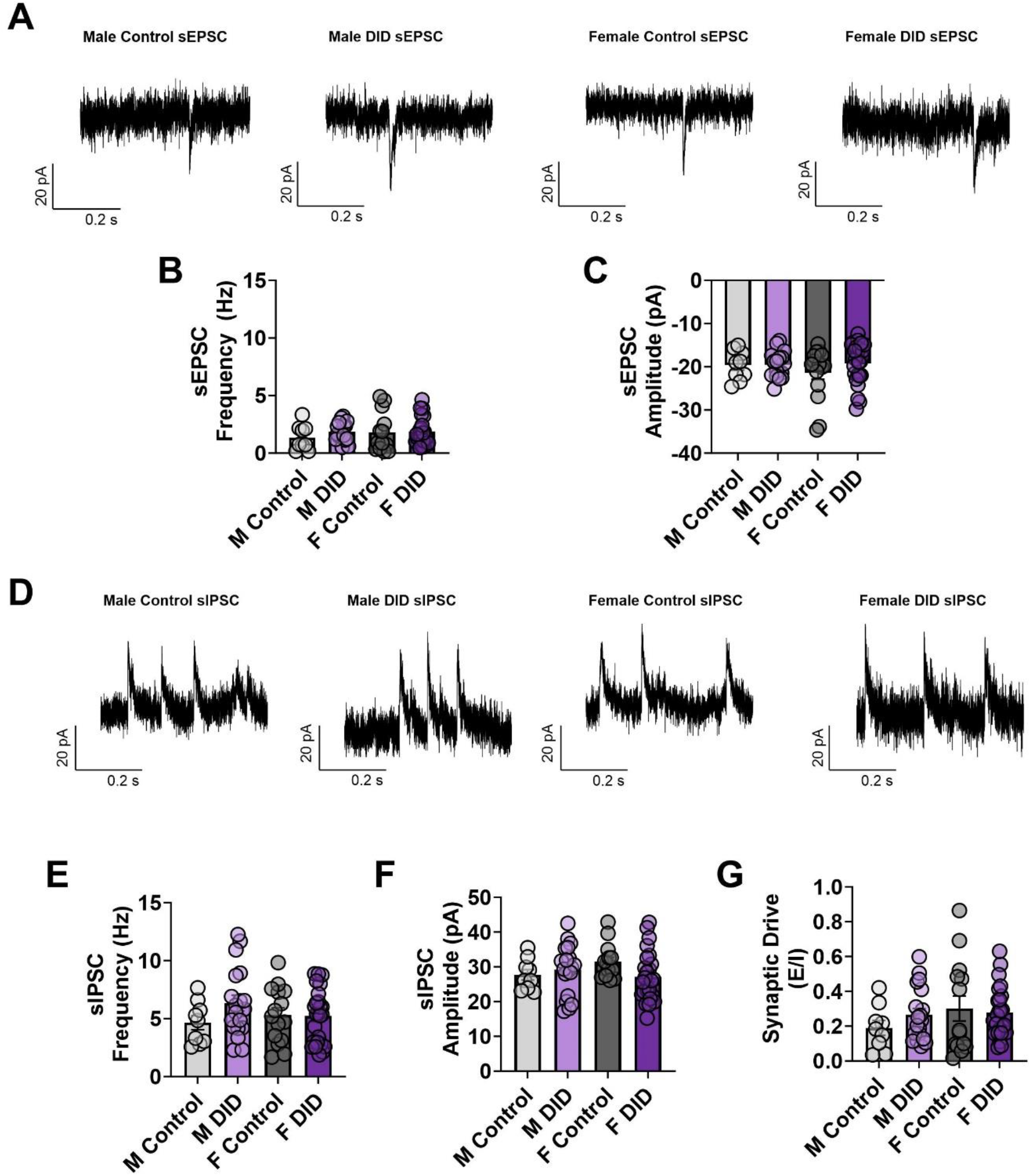
Binge drinking and extended abstinence does not change spontaneous postsynaptic currents onto PL cortex pyramidal neurons. **A)** Representative spontaneous excitatory postsynaptic current (sEPSC) traces. **B)** No change in sEPSC frequency onto pyramidal neurons. **C)** No change in the average amplitude of sEPSC events. **D)** Representative spontaneous inhibitory postsynaptic current (sIPSC) traces**. E)** No change in sIPSC frequency across groups. **F)** No change in average amplitude of sIPSC events. **G)** No change in synaptic drive across groups.

Overall, no changes in frequency or amplitude of sEPSC or sIPSC events were observed in pyramidal neurons across sex or DID condition, indicating that DID does not produce long lasting effects on input events for pyramidal neurons in layer 2/3 of the PL cortex.

### 3.5 Increase in frequency of non-pyramidal spontaneous excitatory post-synaptic currents with modest alcohol exposure

The same experiments outlined above were also performed in non-pyramidal neurons totaling 39 cells from the following groups: female control: n = 7 cells/4 mice, female DID: n = 11 cells/8 mice, male control: 8 cells/4 mice, male DID: 13 cells/8 mice.

In non-pyramidal neurons, there was an effect of DID condition on the frequency of sEPSC events at – 55 mV, where neurons from mice that underwent four weeks of DID had more frequent excitatory events (F_sex_(1, 30) = 0.6385, p = 0.4306; F_DID_(1, 30) = 5.047, p = 0.0322; F_sex x DID_(1, 30) = 0.02382, p = 0.8784; Fig. **5B**). No significant differences across groups were observed in the average amplitude of sEPSC events (F_sex_(1, 35) = 2.653e-007, p = 0.9996; F_DID_(1, 35) = 1.581, p = 0.2170; F_sex x DID_(1, 35) = 0.006943, p = 0.9341; Fig. **5C**).

**Figure. 5.**
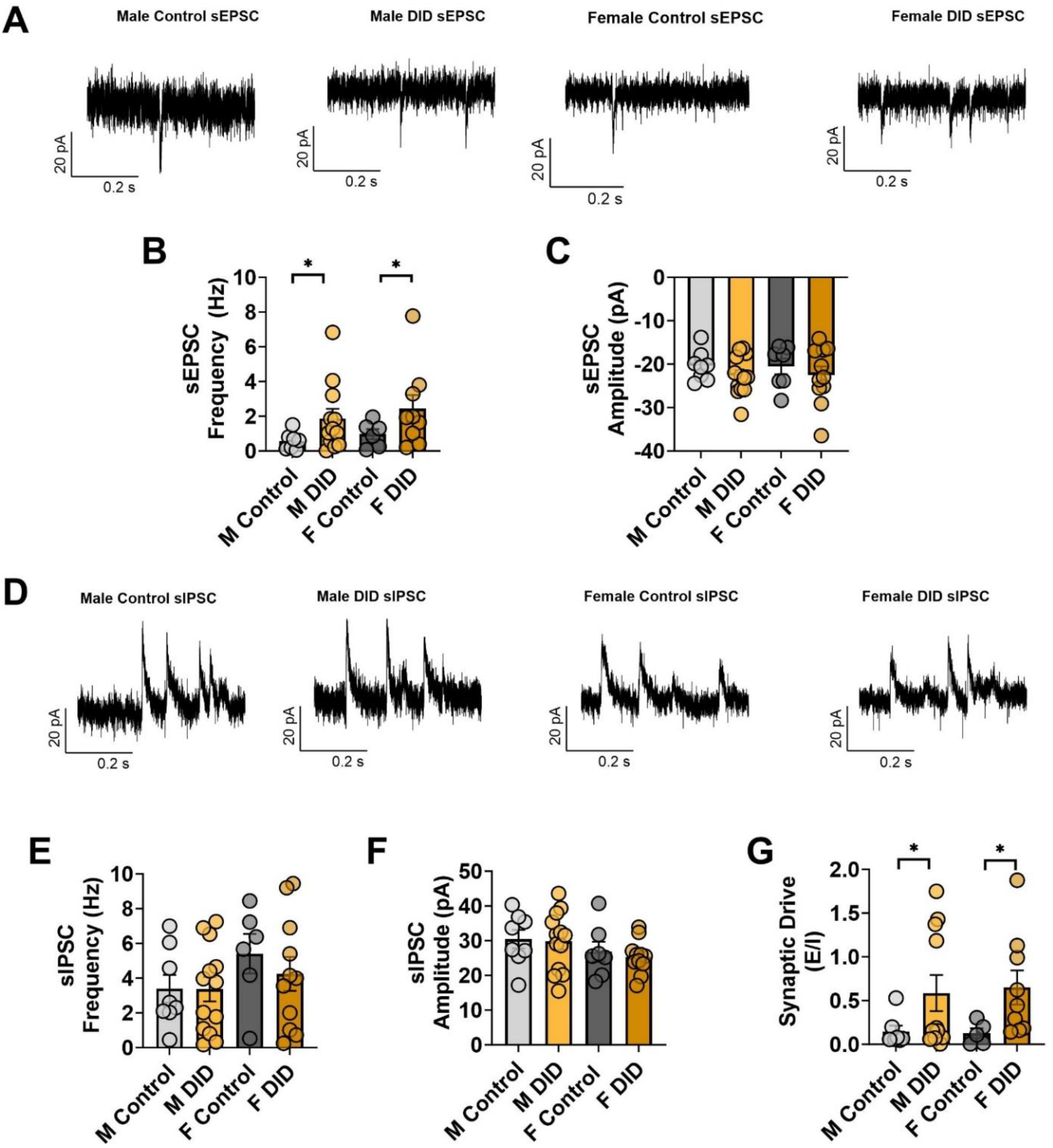
Voluntary binge drinking and extended abstinence induces significant changes in spontaneous glutamatergic, but not GABAergic, inputs onto non-pyramidal neurons. **A)** Representative sEPSC traces. **B)** DID mice show a significant increase in sEPSC frequency onto non-pyramidal neurons. **C)** No change in the average amplitude of sEPSC events. **D)** Representative sIPSC traces. **E)** No change in the frequency of sIPSC events onto non-pyramidal neurons. **F)** No change in average amplitude of sIPSC events. **G)** DID significantly increases the synaptic drive onto non-pyramidal neurons.

No significant differences in non-pyramidal sIPSC frequency or amplitude across sex or DID condition were observed (frequency: F_sex_(1, 34) = 2.439, p = 0.1276; F_DID_(1, 34) = 0.4120, p = 0.5253; F_sex x DID_(1, 34) = 0.3727, p = 0.5456; Fig. **5E**; amplitude: F_sex_(1, 35) = 2825, p = 0.1017; F_DID_(1, 35) = 0.2015, p = 0.6563; F_sex x DID_(1, 35) = 0.02817, p = 0.8677; Fig. **5F**). There was a significant increase in synaptic drive (more excitatory input) onto non-pyramidal neurons in the DID condition (F_sex_(1, 28) = 0.01142, p = 0.9157; F_DID_(1, 28) = 6.191, p = 0.0191; F_sex x DID_(1, 28) = 0.04691, p = 0.8301; Fig. **5G**). The increase in frequency of spontaneous excitatory postsynaptic currents and increase of synaptic drive in non-pyramidal neurons indicates a change in glutamatergic input onto these neurons in the prelimbic region with moderate alcohol consumption.

### 3.6 Individual ethanol consumption levels did not predict changes in postsynaptic transmission onto either cell type in alcohol-exposed individuals

Amongst the DID condition, correlation analysis of each sPSC variable and the mouse’s total EtOH consumption across all four weeks was conducted to determine if the amount of alcohol consumed differentially impacted each measure. There were no correlations in the total alcohol consumed and any of these measurements of spontaneous release (Fig. **6A-E**). Except in the case of sIPSC frequency and total EtOH consumption, where greater EtOH consumption was correlated with lower sIPSC frequency (p = 0.0205; Fig. **6C**). No significant correlations were found, indicating that the total amount of alcohol consumed did not impact the postsynaptic signaling of non-pyramidal neurons (Fig. **6F-J**).

**Figure 6.**
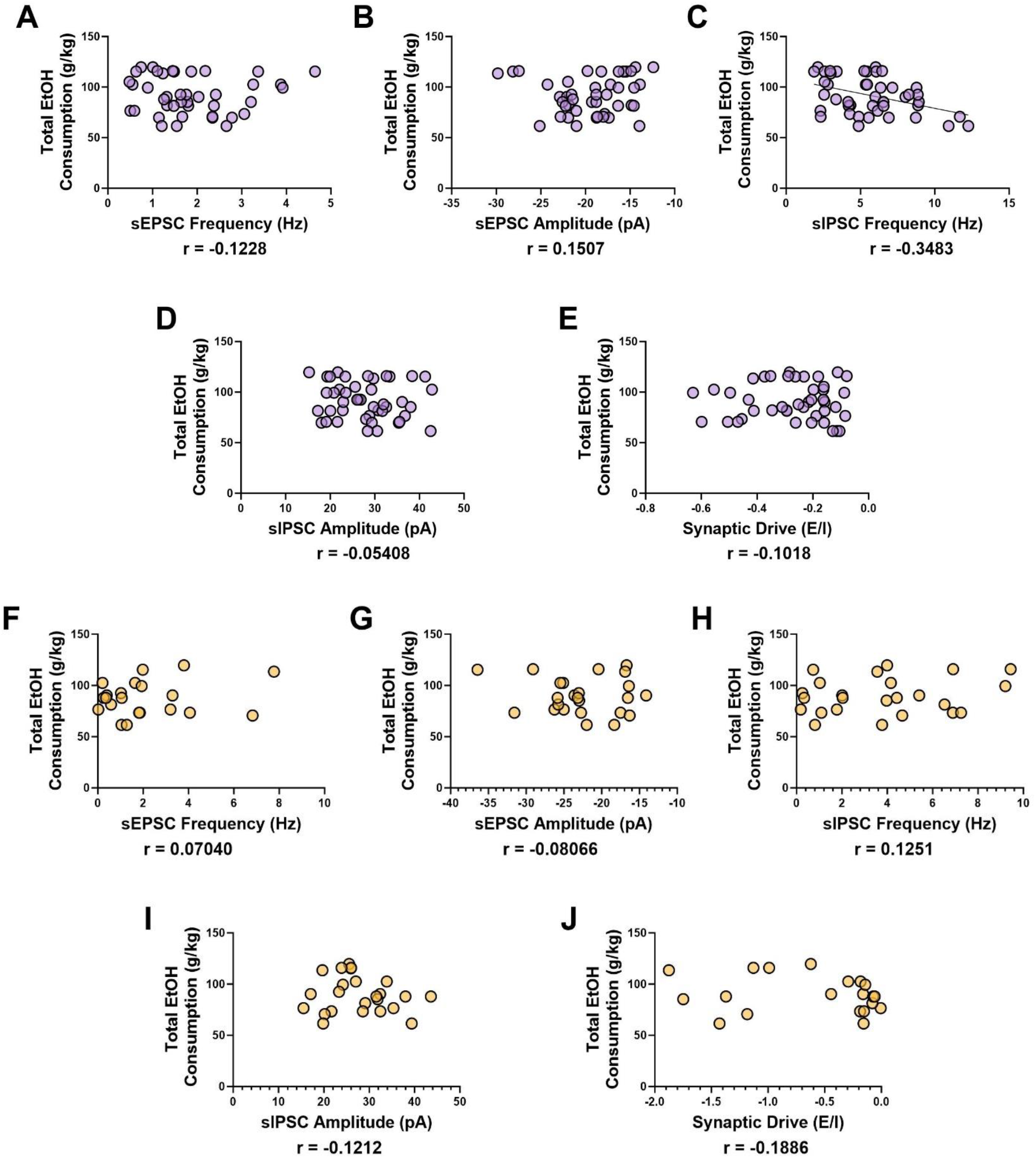
Spontaneous postsynaptic current parameters did not correlate with total ethanol consumed. **A)** Correlation of pyramidal sEPSC frequency and total EtOH consumption not significant (p = 0.4444). **B)** Correlation of pyramidal sEPSC amplitude and total EtOH consumption not significant (p = 0.3347). **C)** Significant negative correlation of pyramidal sIPSC frequency and total EtOH consumption (p = 0.0205). **D)** Pyramidal sIPSC amplitude and total EtOH consumption not significantly correlated (p = 0.5107). **E)** Pyramidal synaptic drive and total EtOH consumption not significantly correlated (p = 0.3653). **F)** Non-pyramidal sEPSC frequency not significantly correlated with total EtOH consumption (p = 0.7617). **G)** Correlation of non-pyramidal sEPSC amplitude and EtOH consumption not significant (p = 0.7079). **H)** Non-pyramidal sIPSC frequency not significantly correlated with total EtOH consumption (p = 0.5602). **I)** No significant correlation between non-pyramidal sIPSC amplitude and total EtOH consumption (p = 0.5726). **J)** No significant correlation between non-pyramidal synaptic drive and total EtOH consumption (p = 0.4258).

## 4. DISCUSSION

Here we present, to our knowledge, the first evidence that moderate voluntary binge alcohol consumption induces changes in intrinsic excitability and the excitation/inhibition balance after a prolonged period of abstinence (∼6 months) within the prefrontal cortex, a region known to be involved in the pathology of AD. Specifically, while alcohol exposure did not alter sIPSCs or sEPSCs onto layer 2/3 pyramidal neurons of the PL cortex, it did induce changes in intrinsic excitability in this population, where alcohol consumption lead to a hyperpolarization of the RMP of pyramidal neurons, while also altering the AP threshold, in both male and female mice. While this effect is seen in both males and females, some sex differences were detected in the rheobase and VI at RMP. Male pyramidal neurons fire more readily than female pyramidal neurons; this effect is predominately driven by sex differences in the control animals, whereas DID seems to neutralize this effect. This interaction between sex and alcohol-induced changes is important to further investigate, given the notable sex difference in prevalence of AD (Seemiller et al., 2024).

In a study with the same DID paradigm, neurons in layer 2/3 of the PL cortex were found to have increased intrinsic excitability following DID 24 h following alcohol (Dao et al., 2021). More specifically, the increase in pyramidal intrinsic excitability was defined by a decrease in rheobase and increase in the number of APs fired in VI, with no changes in RMP or AP threshold (Dao et al., 2021). Here we find that at protracted timepoints (approximately 6 months following DID), female pyramidal neurons also show increased AP firing during VI, while males have undergone a shift towards decreased firing, highlighting potential compensatory homeostatic regulation may occur following DID to attenuate the initial increase in pyramidal intrinsic excitability. The mechanisms underlying this homeostatic regulation are unknown but may include changes in the expression of voltage-gated sodium channels. Importantly, control groups differ in their intrinsic excitability measures across our paper and Dao et al., indicating a basal effect of aging on pyramidal excitability, highlighting that more work is needed to understand the effect of healthy natural aging of the prefrontal cortex.

Findings in non-pyramidal neurons further support the hypothesis that a homeostatic regulation of signaling in the PL cortex occurs in the months following binge drinking. In non-pyramidal neurons, the changes are seen in both intrinsic firing properties and presynaptic excitatory signaling onto the neurons. Some modulation of intrinsic excitability may be occurring, as we observed that non-pyramidal neurons from DID mice had less negative AP thresholds at –70 mV. Holding the cells at –70 mV allows for standardization across membrane potential, and on average the –70 mV was approximately 5 mV more hyperpolarized than RMP, suggesting fewer voltage-gated channels (Na^+^, Ca^+^, Cl^-^) are open.

Spontaneous signaling onto non-pyramidal neurons was significantly altered with binge drinking. The more than two-fold increase in sEPSC frequency and corresponding increase in synaptic drive indicate an upstream increase in glutamatergic transmission onto GABAergic neurons. Additionally, DID increases the expression of glutamatergic receptors in the prefrontal cortex (Szumlinkski et al., 2023), which complements and provides a potential mechanism underlying findings here. Exposure to alcohol in the immediate term is known to decrease interneuron excitability in the PL cortex and decrease sEPSC frequency in the somatostatin subtype (Thompson et al., 2019; Dao et al., 2021). The hippocampal input to the PFC is a strong candidate for this change (O’Donnell & Grace, 1995; Ruggiero et al., 2021; Song et al., 2022).

Seemingly, the inputs to pyramidal and non-pyramidal neurons in the PL cortex are either differentially impacted following homeostasis after alcohol consumption or the increase of input to the inhibitory interneurons neutralizes an increase of excitation. Layer 2/3 of the PL cortex has been implicated as a critical region for maintaining proper cortical E/I balance, as it receives the most innervation from subcortical regions (Fagerholm et al., 2016). The increased excitatory input could result in greater synchrony of inhibition, which if seen in humans following prolonged alcohol use, would worsen E/I balance in AD. Finally, because the magnitude of increase is over twice the amount of excitation, excitotoxicity may occur in non-pyramidal neurons following binge alcohol consumption and prolonged abstinence. In this case, alcohol consumption would significantly worsen AD E/I imbalance as there is already a significant loss in a subtype of non-pyramidal somatostatin expressing neurons (Davies et al., 1980; Burgos-Ramos et al., 2008).

## 5. CONCLUSION

Findings here illuminate that homeostatic regulation occurs in the PL cortex in the months following binge alcohol consumption, resulting in a potential decrease in pyramidal neuron intrinsic excitability and a two-fold increase in sEPSC frequency and synaptic drive onto non-pyramidal neurons. Work here suggests that alcohol could be a long-term modulator of E/I balance, creating a neuronal environment similar to that of early AD.

## ABBREVIATIONS

aCSF: artificial cerebrospinal fluid
AP: action potential
AUD: alcohol use disorder
ADRD: Alzheimer’s Disease and Related Dementias
DID: Drinking-in-the-Dark
E/I: excitatory/inhibitory
EtOH: ethanol
fAHP: fast afterhyperpolarization
NMDG: N-methyl-D-glucamine
PL: prelimbic
RMP: resting membrane potential
sEPSC: spontaneous excitatory postsynaptic current
sIPSC: spontaneous inhibitory postsynaptic current
VI: current injection

## 6. ACKNOWLEDGEMENTS

Funding: R01AA029403 to NAC, F31AA030455 to ARS, R01AG072513 to EAP, and The Huck Institutes of the Life Sciences Endowment Funds to NAC

## CRediT authorship contribution statement

**Grace C. Smith:** Conceptualization, Data curation, Formal analysis, Investigation, Methodology, Software, Writing – original draft, Writing – review & editing. **Keith R. Griffith:** Investigation, Data curation, Formal analysis, Software, Visualization, Writing – review & editing. **Avery R. Sicher:** Investigation, Data curation, Formal analysis, Writing – review & editing. **Dakota F. Brockway:** Investigation, Data curation, Writing – review & editing. **Elizabeth A. Proctor:** Formal analysis, Funding acquisition, Software, Writing – original draft, Writing – review & editing. **Nicole A. Crowley:** Conceptualization, Formal analysis, Funding acquisition, Investigation, Methodology, Project administration, Resources, Supervision, Writing – original draft, Writing – review & editing.

